# Cholinergic feedback for context-specific modulation of sensory representations

**DOI:** 10.1101/2024.09.10.612274

**Authors:** Bin Yu, Yuxuan Yue, Chi Ren, Rui Yun, Byungkook Lim, Takaki Komiyama

**Affiliations:** Department of Neurobiology, Department of Neurosciences, Center for Neural Circuits and Behavior, Kavli Institute for Brain and Mind, and Halıcıoğlu Data Science Institute, University of California San Diego, La Jolla, CA, USA; Department of Electrical and Computer Engineering, University of California San Diego, La Jolla, CA, USA

## Abstract

The brain’s ability to prioritize behaviorally relevant sensory information is crucial for adaptive behavior, yet the underlying mechanisms remain unclear. Here, we investigated the role of basal forebrain cholinergic neurons in modulating olfactory bulb (OB) circuits in mice.

Calcium imaging of cholinergic feedback axons in OB revealed that their activity is strongly correlated with orofacial movements, with little responses to passively experienced odor stimuli. However, when mice engaged in an odor discrimination task, OB cholinergic axons rapidly shifted their response patterns from movement-correlated activity to odor-aligned responses.

Notably, these odor responses during olfactory task engagement were absent in cholinergic axons projecting to the dorsal cortex. The level of odor responses correlated with task performance. Inactivation of OB-projecting cholinergic neurons during task engagement impaired performance and reduced odor responses in OB granule cells. Thus, the cholinergic system dynamically modulates sensory processing in a modality-specific and context-dependent manner, providing a mechanism for a flexible and adaptive sensory prioritization.

## Main Text

Animals are constantly bombarded by a plethora of incoming information, making the brain’s ability to prioritize behaviorally relevant information critical for adaptive and flexible behavior. Cholinergic neurons in the basal forebrain (BF), a principal source of acetylcholine in the brain, are considered central to this brain function. Cholinergic neurons project extensively to many brain areas and modulate circuit functions and plasticity (*1–3*). In sensory regions, acetylcholine can enhance the gain of sensory neurons (*4–9*), leading to the hypothesis that cholinergic signaling contributes to sensory attention which amplifies the representations of behaviorally relevant stimuli (*10–14*). Moreover, the decline in cognitive functions, such as selective attention and cognitive flexibility, particularly evident in Alzheimer’s and Parkinson’s diseases, is associated with the progressive loss of cholinergic neurons (*15, 16*). Notably, deep brain stimulation of the basal forebrain is being explored as a therapeutic option for dementia, showing promise in alleviating cognitive symptoms in some patients with Alzheimer’s and Parkinson’s-related dementia (*17, 18*).

However, recent studies directly recording the activity of BF cholinergic neurons challenge the notion that cholinergic neurons regulate selective attention by selectively enhancing the processing of relevant information. For instance, a large number of studies have found that cholinergic activity correlates strongly with pupil size, locomotion, and facial movement, suggesting that it might be more closely related to general arousal rather than selective attention (*19–26*). For cholinergic neurons to control selective attention, their functional properties would need to be dynamic, adapting to behavioral demands. Here we explored this idea in mice during odor-guided behavior.

## Behavioral context switches the tuning of OB cholinergic axons

To investigate the role of cholinergic activity in olfactory processing, we used a two-alternative choice odor discrimination task in head-fixed mice. Water-restricted mice were trained under head fixation to discriminate between different binary mixture ratios (60:40, 53:47, 47:53, and 40:60) of heptanal and ethyl tiglate and indicate their choice by licking either the left or right lickport. The correct choices were left for 60:40 and 53:47 and right for 47:53 and 40:60, with water as the reward. A 2-second sound cue preceded a 4-second odor presentation, and mice were required to respond within 2 seconds after the odor offset (**Fig. 1A,B**). A separate cohort of mice underwent the passive paradigm, in which mice experienced the same odors in the same temporal structure without ever engaging in a task. We expressed axon-GCaMP6s (*27*) in BF cholinergic neurons by injecting Cre-dependent AAV in the BF of ChAT-Cre mice and imaged cholinergic axon activity in the olfactory bulb (OB) during the task and passive paradigms by two-photon microscopy (**Fig. 1C,D**).

**Fig. 1.**
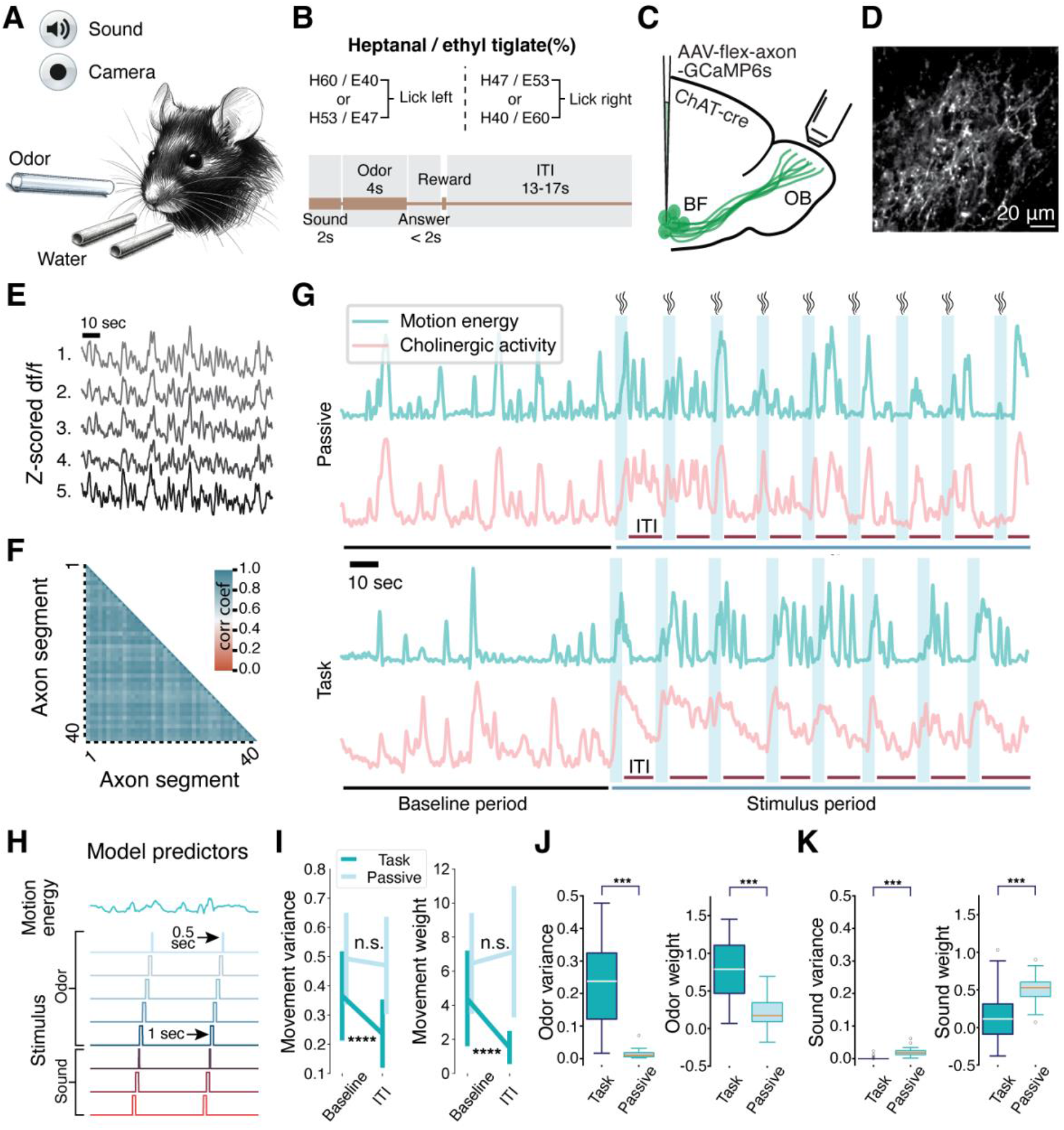
Behavioral context switches the tuning of OB cholinergic axons. (**A**) Schematic of the experimental setup. (**B**) Task structure. Mice discriminated between different ratios of heptanal and ethyl tiglate. A 2-second sound cue preceded a 4-second odor presentation. Mice indicated their choice by licking left or right within 2 seconds after odor offset. Correct choices were rewarded by water. (**C**) Viral strategy for expressing GCaMP6s in basal forebrain cholinergic neurons. Their axons were imaged in the OB. (**D**) Representative two-photon image of cholinergic axons expressing GCaMP6s in the OB. Scale bar: 20 μm. (**E**) Example time series of GCaMP6s fluorescence from 5 axon segments imaged simultaneously, showing high correlation. (**F**) Correlation matrix of GCaMP6s signals from all 41 axon segments imaged simultaneously, demonstrating high synchrony across segments. (**G**) Representative time series of orofacial motion energy (blue) and cholinergic axon activity (red) during passive odor exposure (top) and task engagement (bottom). Odor presentations are indicated by light blue shading. (**H**) Ridge regression predictors for orofacial motion energy, odor stimuli (1st - 4th sec and a 0.5 second delay period), and sound cue (1st, 2nd sec and a 0.3 second delay period) for the generalized linear model predicting cholinergic axon activity. (**I**) The variance (left) and weight (right) for the motion energy predictor between baseline and inter-trial interval (ITI) periods for task and passive conditions. Wilcoxon signed-rank test, *p* = 1.13 × 10^-6^ and *p* = 1.06 × 10^-9^ for task variance and weight, *p* = 0.57 and *p* = 0.062 for passive variance and weight, mean ± std. n = 1157 and 833 axon segments from 55 and 32 imaging fields from 6 and 7 animals for task and passive respectively. (**J**) Odor variance (left) and weight (right) between task and passive conditions. Wilcoxon rank-sum test, *p* = 4.89 × 10^-14^ and *p* = 6.9 × 10^-11^ for variance and weight). (**K**) Sound variance (left) and weight (right) between task and passive conditions. Wilcoxon rank-sum test, *p* = 1.16 × 10^-13^ and *p* = 1.23 × 10^-8^ for variance and weight. Statistical significance: *p < 0.05, **p < 0.01, ***p < 0.001, n.s.: p > 0.05.

We observed that all segments of cholinergic axons within each field of view were highly temporally correlated (**Fig. 1E,F**). Thus, we used the average activity across segments for the subsequent analysis. The animals’ orofacial movements were quantified by calculating the frame-by-frame differences in the recorded facial videos, referred to as motion energy. Before the first odorant delivery, we recorded the spontaneous activity of cholinergic axons in the OB for approximately 130 seconds, a period designated as the baseline. Following this, the period after the onset of the first odorant delivery was defined as the stimulus period.

In the baseline period for both task and passive animals, we found that cholinergic axons showed substantial spontaneous activity that was strongly correlated with the animals’ orofacial movements—large orofacial movements coincided with large cholinergic activity. In the passive condition, this strong correlation with orofacial movements continued in the stimulus period, and there was little response time-locked to the odorant stimuli. In stark contrast, when the animals were engaged in the odor discrimination task, the cholinergic axons quickly shifted their functional properties and began exhibiting strong, phasic responses to odor stimuli. In parallel, movement-correlated responses became weaker when mice engaged in the olfactory task (**Fig. 1G**).

To quantify the degree of movement and sensory responses, we applied generalized linear model analysis with the odor, motion energy, and sound as predictors (**Fig. 1H**, see *Methods*). For movement responses, we chose the periods without sensory stimuli, namely the baseline and inter-trial interval (ITI) periods (**Fig. 1G**). In the passive condition, the strong movement responses were not affected by odor presentation, shown by equivalent variance and weights for movements in baseline and ITI periods. In contrast, task engagement led to a notable decrease in movement variance and weight, indicating a reduction in movement responses (**Fig. 1I**).

Mirroring these reduced movement responses, the variance and weights of odor predictors were substantially higher in the task condition than passive, highlighting that cholinergic axons respond robustly to olfactory stimuli during task engagement (**Fig. 1J**). Responses to the sound cue were weak in both task and passive conditions (**Fig. 1K**). These quantifications demonstrate that the cholinergic axons respond strongly to orofacial motion during the baseline period and passive odor experience, and their tuning rapidly shifts to respond predominantly to odor stimuli during task engagement, highlighting their profound dependence on behavioral context.

## Odor response of cholinergic axons in OB reflects task engagement level and performance during discrimination task

The results above indicate that OB cholinergic axons respond to odor stimuli specifically during task engagement. We next investigated whether the degree of these odor responses was further modulated based on the level of task engagement. To address this, we adopted an analytical method that identifies trial blocks within individual sessions with different behavioral states.

This method, known as the hidden Markov model with generalized linear model observations (GLM-HMM), takes task variables as predictors, including the animal’s previous trial choice, the current stimulus, and the current trial’s choice, to decompose the animal’s behavioral states (*28*).

By fitting GLM-HMM to the behavioral data during task engagement, we uncovered three distinct behavioral states across sessions. In one state, mice chose either the left or right side based on the odor, leading to a high reward rate in this state, which we refer to as the engaged state. In the other two states, the mice were biased to one choice regardless of the presented odor, leading to ∼50% reward rate. We refer to these states as biased states. In many imaging sessions, mice transitioned between these states, while mice remained in only one state in some sessions. Approximately half of all trials were classified as the engaged state, while the remainder fell into the biased states (**Fig. 2A-E**).

**Fig. 2.**
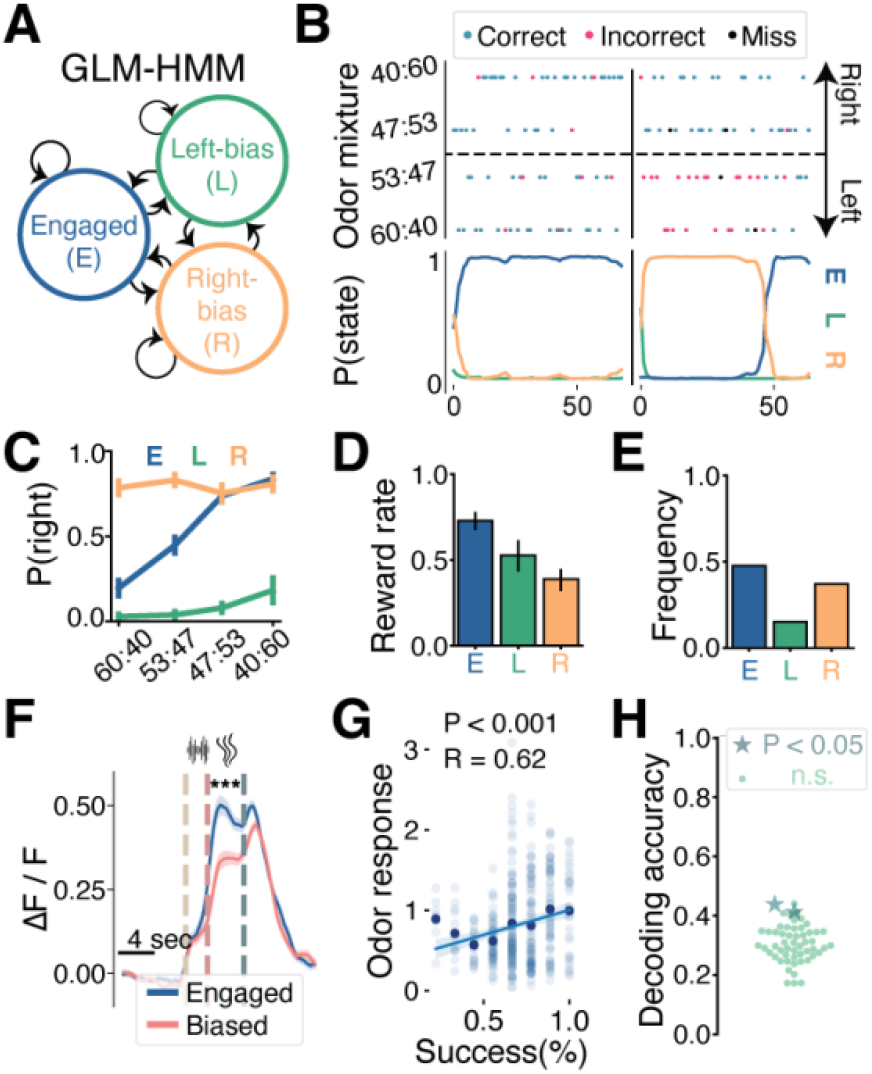
Odor response of OB cholinergic axons reflects the level of task engagement. (**A**) Schematic of the Generalized Linear Model-Hidden Markov Model (GLM-HMM) used to identify behavioral states. (**B**) Choices (top) and corresponding state probabilities (bottom) identified by GLM-HMM from two example sessions. (**C**) Probability of right choice for each odor mixture in the three behavioral states. Mean ± SEM. (**D**) Reward rates in each behavioral state. Mean ± SEM. (**E**) Frequency of each behavioral state across all sessions. N = 55 sessions from 6 mice. (**F**) Trial-averaged cholinergic axon responses to odor stimuli in engaged (blue) and biased (red) states. Shades represent SEM (Wilcoxon rank-sum test, *p* = 4.89 × 10^-7^). N = 444 (engaged) and 537 (biased) trials from 19 FOVs from 6 mice. (**G**) Correlation between odor response amplitude and behavioral success rate. Each dot represents a sliding window of 9 trials. Linear regression, *p* < 0.001, R = 0.62. (**H**) Decoding accuracy of odor mixture identity from single-trial responses of individual cholinergic axon segments. Each dot represents one imaging session. Stars indicate sessions with significant decoding (permutation test, *p* = 0.019 and *p* = 0.03). Cholinergic axons do not consistently encode odor identity. Statistical significance: *p < 0.05, **p < 0.01, ***p < 0.001, n.s.: p > 0.05.

We then examined odor responses of OB cholinergic axons in sessions that contained both engaged state and at least one of the biased states. Our analysis revealed that cholinergic axons responded more strongly to odors in the engaged state than in the biased states (**Fig. 2F**). We further examined trials within engaged state. Using a sliding window of nine trials, we analyzed the odor response amplitude against the animal’s success rate, which uncovered that higher success rates were associated with stronger odor responses (**Fig. 2G**). Thus, the amplitude of odor responses of cholinergic axons in the task condition is graded such that more engaged and better performing trials were associated with stronger odor responses.

We next asked whether cholinergic axons’ phasic odor responses encoded the odor identity. We tested this by utilizing a support vector machine (SVM) to decode odor mixture identity on individual trials based on the odor response of single axons (we did not average across axonal segments for this analysis only). The decoding accuracy was at chance level in all but 2 sessions, indicating that these cholinergic axons did not strongly encode odor identity (**Fig. 2H**).

In summary, we found that cholinergic axons show stronger odor responses when mice are more engaged in odor discrimination, suggesting that cholinergic signaling modulates OB odor processing based on the degree of task engagement. The strength of these odor responses also correlated with task performance.

## Region-Specific Modulation and Targeting of OB-Projecting Cholinergic Neurons

Cholinergic neurons as a whole project to many brain regions, and it is unclear whether cholinergic projections to different brain regions can be differentially modulated. We hypothesized that the observed shift in the functional tuning of OB cholinergic axons during the olfactory task is region-specific. To test this hypothesis, we utilized a large chronic cranial window (*29*) that exposed most of the dorsal cortex and performed two-photon calcium imaging of cholinergic axons in several cortical regions during the same olfactory discrimination task (**Fig. 3A**). We focused on the primary motor cortex (M1), primary somatosensory cortex (S1), and visual areas, which include the primary visual cortex and nearby higher-order visual regions (**Fig. 3B**).

**Fig. 3.**
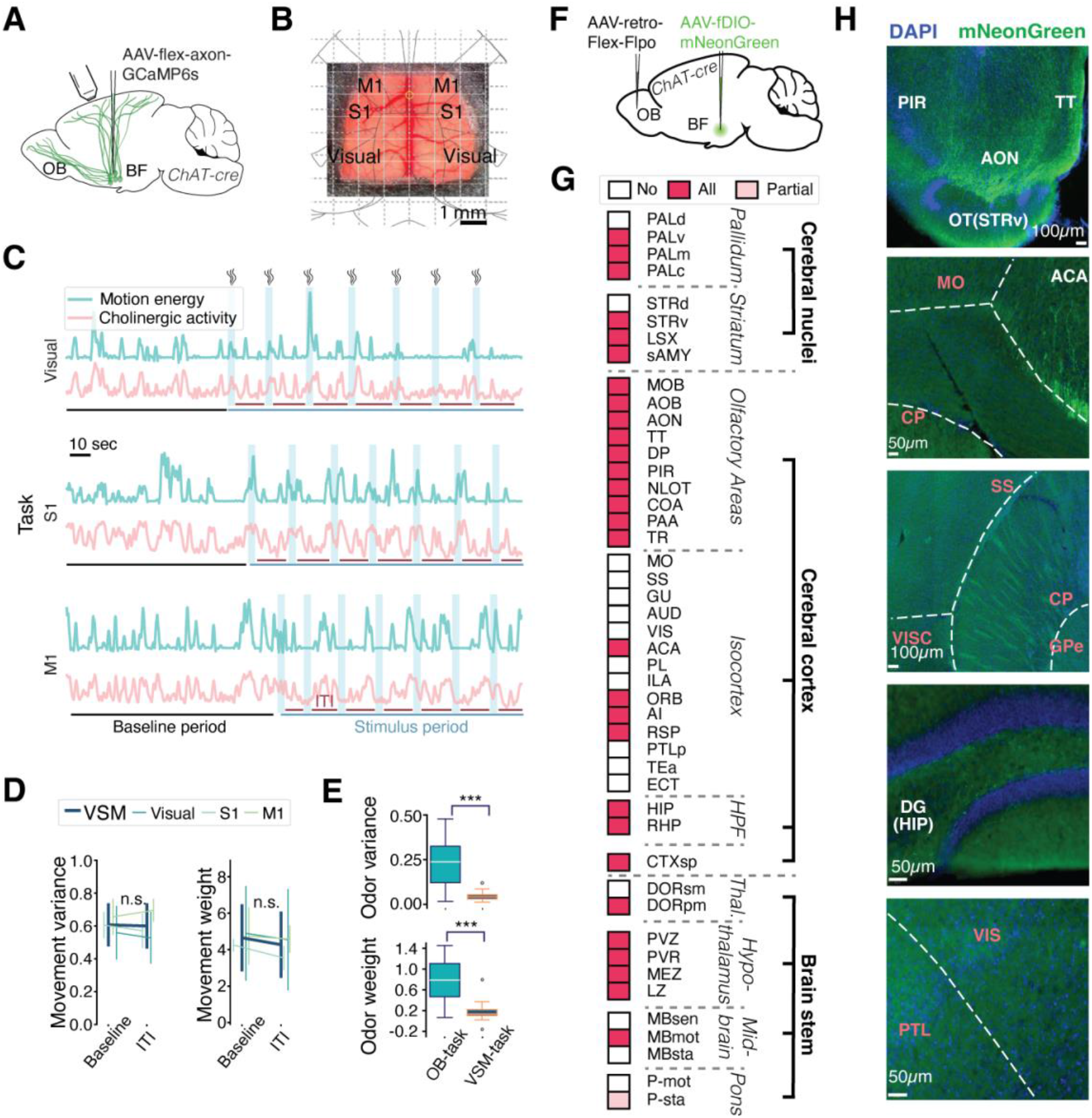
Area-specificity of cholinergic modulation. (**A**) Schematic of imaging cholinergic axons projecting to the dorsal cortex. (**B**) Large chronic cranial window exposing multiple cortical regions. (**C**) Representative time series of orofacial motion energy (blue) and cholinergic axon activity (red) in visual, primary somatosensory (S1), and primary motor (M1) cortices during task engagement. Light blue shading indicates odor presentation periods. (**D**) The variance and weight for orofacial motion energy in baseline and inter-trial interval (ITI) periods for visual, S1, and M1 cortices (VSM is mean across the three areas). Mixed effects model, *p* = 0.887 and *p* = 0.602 for variance and weight, n = 14 FOVs from 4 mice, mean ± std. (**E**) Odor variance and weight in OB and VSM during task engagement. Wilcoxon rank-sum test, *p* = 1.27 × 10^-7^ and *p* = 9.15 × 10^-7^ for variance and weight. (**F**) Viral strategy for tracing collateral projections of OB- projecting cholinergic neurons. (**G**) Mapping of collateral projections from OB-projecting cholinergic neurons. The listed areas are targets of basal forebrain cholinergic neurons (*30*). OB- projecting cholinergic neurons target a subset of these areas. Partial: axons observed in at least one mouse but not all, All: axons observed in all mice. N = 3 mice. (**H**) Representative images of mNeonGreen-labeled axons (green) of OB-projecting cholinergic neurons in various brain regions. DAPI in blue. PIR: piriform cortex, TT: taenia tecta, AON: anterior olfactory nucleus, OT(STRv): olfactory tubercle (ventral striatum), ACA: anterior cingulate area, MO: motor cortex, SS: somatosensory cortex, CP: caudoputamen, VISC: visceral area, GPe: globus pallidus externa, DG(HIP): dentate gyrus (hippocampus), VIS: visual cortex, PTL: posterior parietal association areas. Statistical significance: *p < 0.05, **p < 0.01, ***p < 0.001, n.s.: p > 0.05.

In all these regions, during the baseline period, cholinergic axon activity strongly correlated with the facial motion energy, similarly to OB cholinergic axons. In contrast to OB, during task engagement, the cortical cholinergic axons did not respond to odor stimuli and instead continued to respond during orofacial movements (**Fig. 3C**). We quantified this observation using the same regression analysis as in Fig. 1, and we found that cholinergic axon activity in these cortical regions were similarly correlated with the animals’ orofacial movements during baseline and ITI periods (**Fig. 3D**). The odor responses in these cortical regions were substantially lower than those observed in the OB during task engagement (**Fig. 3E**). These findings indicate that the shift in tuning towards odor stimuli during olfactory task engagement is specific to the OB-projecting cholinergic neurons and not observed in the cholinergic projections to M1, S1, and visual areas.

The results above suggest that the OB-projecting cholinergic neurons are distinct from the cholinergic neurons projecting to the dorsal cortex. We next sought to determine whether the OB-projecting cholinergic neurons also project to other regions. To achieve this goal, we injected retrograde AAV encoding Cre-dependent Flp into the OB and AAV encoding Flp-dependent mNeonGreen into the basal forebrain of ChAT-cre mice. Brain tissues were processed two to three weeks post-injection, images were registered to the Allen reference atlas, and the areas with labeled axons were identified. We focused our analysis on the regions previously shown to receive projections from cholinergic neurons in the basal forebrain (*30*). We observed labeled axons in roughly half of these regions in the hemisphere ipsilateral to the injection site. Notably, many of the regions that received collateral projections from the OB-projecting cholinergic neurons were closely associated with the olfactory system, such as hippocampus, hypothalamus and frontal cortex. No projections were detected in the primary motor and sensory regions (**Fig. 3F**). This finding suggests that the OB-projecting cholinergic neurons constitute a distinct subgroup primarily targeting the olfactory pathway.

Our findings indicate that cholinergic neurons projecting to the olfactory pathway are a distinct population from those projecting to the dorsal cortex. The context-dependent modulation of cholinergic axons during the olfactory discrimination task is predominantly specific to this OB-projecting population and does not occur in other sensory and motor regions.

## OB-projecting cholinergic neurons modulate OB neural activity and contribute to olfactory behavior

Next, we aimed to examine the impact of OB-projecting cholinergic neurons on OB activity and animal behavior. Previous studies have identified the granule cells, local inhibitory neurons and the most abundant neural population in the OB, as the main target of cholinergic feedback (*31, 32*). Therefore, we focused our analysis on granule cells. To image granule cell activity, we injected AAV encoding GCaMP6f. Five weeks after the injection, we imaged granule cell populations in mice performing the olfactory discrimination task (n = 4 mice) and other mice in the passive odor exposure paradigm (n = 4 mice), similar to imaging of cholinergic axons (**Fig. 4A**, **Fig. 1A-B**). We identified granule cell-odor pairs that show odor-excited and odor-suppressed responses (**Fig. 4B**, see *Methods*). We found that similar fractions of cell-odor pairs showed odor responses in the task and passive conditions, but the amplitude of their responses was significantly larger in the task condition for both excited and suppressed responses (**Fig. 4C, D**).

**Fig. 4.**
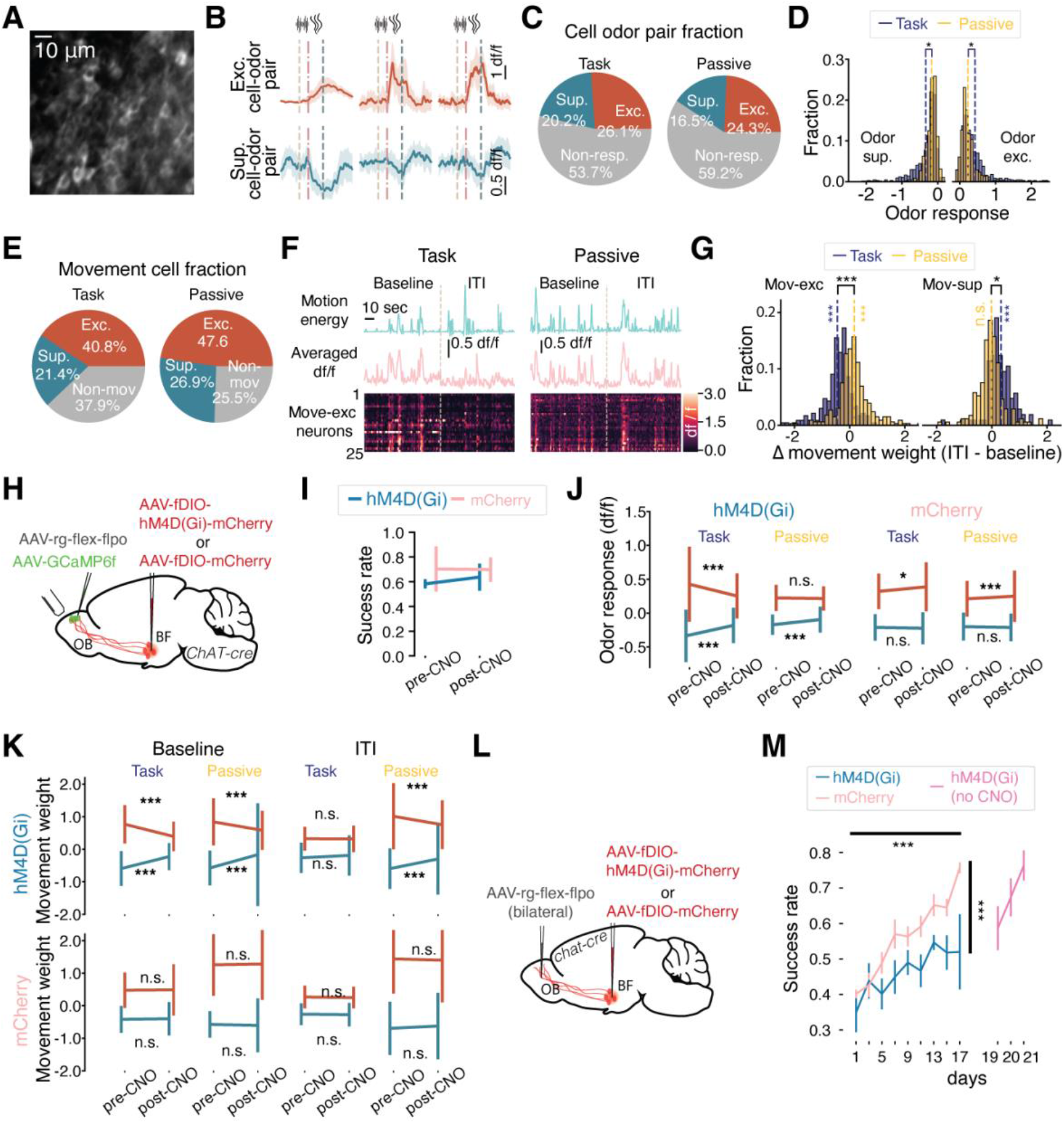
OB-projecting cholinergic neurons modulate OB neural activity and contribute to olfactory behavior. (**A**) Representative two-photon image of OB granule cells expressing GCaMP6f. (**B**) Example odor responses of excited (top) and suppressed (bottom) granule cells. Mean ± SEM. (**C**) Fractions of excited (Exc), suppressed (Sup), and non-responsive (Non-resp) granule cell-odor pairs of total cell-odor pairs during task engagement and passive exposure. n = 1011 and 1164 granule cells from 8 animals for both task and passive respectively. In total, there are 4044 and 4656 possible cell-odor pairs. (**D**) Distribution of odor response amplitudes for suppressed and excited cell-odor pairs during task and passive conditions. The amplitude of odor responses was significantly larger in the task condition (mixed effects model, *p* = 0.03 and *p* = 0.019 for excited and suppressed cell-odor pair). (**E**) Fractions of movement-excited (Exc), movement-suppressed (Sup), and non-movement-responsive (Non-mov) neurons of total granule cells during task and passive conditions. (**F**) Representative time series of orofacial motion energy (top) and averaged movement-excited granule cell activity (middle) during task and passive conditions. Bottom: activity heatmap of 25 movement-excited granule cells sorted by movement weight. Note the decrease in movement responses with task engagement. (**G**) Distribution of changes in movement weights between ITI and baseline periods for movement- excited (Mov-exc) and movement-suppressed (Mov-sup) neurons during task and passive conditions. Mixed effects model, *p* < 0.001 and *p* < 0.001 for excited cell-odor pairs of task and passive conditions compare to 0; *p* < 0.001 and *p* = 0.638 for suppressed cell-odor pairs of task and passive conditions compare to 0; *p* < 0.001 and *p* = 0.019 for excited and suppressed cell- odor pairs between task and passive conditions. (**H**) Viral strategy for expressing hM4D(Gi) or mCherry in OB-projecting cholinergic neurons and GCaMP6f in granule cells. (**I**) Behavioral success rates before and after CNO administration in hM4D(Gi) and mCherry control groups. This unilateral manipulation did not affect behavior. (**J**) Odor response amplitudes of granule cells before and after CNO administration in task and passive conditions for hM4D(Gi) and mCherry groups. Mixed effects model, *p* < 0.001 and *p* = 0.152 for hM4D(Gi) odor-excited cell- odor pairs in task and passive conditions, *p* < 0.001 for hM4D(Gi) odor-suppressed cell-odor pairs in both task and passive conditions, *p* = 0.013 and *p* < 0.001 for mCherry odor-excited cell- odor pairs in task and passive conditions, *p* = 0.061 and *p* = 0.43 for mCherry odor-suppressed cell-odor pairs in task and passive conditions. n = 695 and 703 from 4 animals each for hM4D(Gi) excited cell-odor pairs for task and passive; n = 573 and 486 from 4 animals each for hM4D(Gi) suppressed cell-odor pairs for task and passive; n = 427 and 359 from 4 animals each for mCherry excited cell-odor pairs for task and passive; n = 245 and 200 from 4 animals each for mCherry suppressed cell-odor pairs for task and passive. (**K**) Movement weights of granule cells during baseline and ITI periods, before and after CNO administration, in task and passive conditions for hM4D(Gi) and mCherry groups. Mixed effects model comparing pre- and post- CNO, *\*p* < 0.001 for both movement-excited and movement-suppressed granule cell baseline weight in both task and passive conditions of hM4D(Gi) group, *p =* 0.762 and *p <* 0.001 for movement-excited granule cell ITI weight in task and passive conditions of hM4D(Gi) group. *p =* 0.17 and *p <* 0.001 for movement-suppressed granule cell ITI weight in task and passive conditions of hM4D(Gi) group. *p =* 0.78 and *p =* 0.68 for movement-excited granule cell baseline weight in task and passive conditions of mCherry group. *p =* 0.67 and *p =* 0.64 for movement-suppressed granule cell baseline weight in task and passive conditions of mCherry group. *p =* 0.38 and *p =* 0.50 for movement-excited granule cell ITI weight in task and passive conditions of mCherry group. *p =* 0.46 and *p =* 0.14 for movement-suppressed granule cell ITI weight in task and passive conditions of mCherry group. n = 232 and 332 from 4 animals each for hM4D(Gi) movement-excited cells for task and passive respectively; n = 105 and 172 from 4 animals each for hM4D(Gi) movement-suppressed cells for task and passive; n = 180 and 222 from 4 animals each for mCherry movement-excited cells for task and passive; n = 111 and 141 from 4 animals each for mCherry movement-suppressed cells for task and passive. (**L**) Viral strategy for bilateral expression of hM4D(Gi) or mCherry in OB-projecting cholinergic neurons. (**M**) Behavioral success rates over training days for hM4D(Gi) and mCherry groups. Two-way ANOVA, group: *p =* 4.71 × 10^-6^, day: *p =* 2.26 × 10^-7^, mean ± sem, n = 5 mice for each group. Data are presented as mean ± std unless otherwise noted. Statistical significance: *p < 0.05, **p < 0.01, ***p < 0.001, n.s.: p > 0.05.

Next we examined whether granule cells exhibit activity correlating with orofacial movements. Quantifying movement modulations of individual granule cells using GLM with orofacial motion energy as a predictor, we identified movement-excited and movement-suppressed neurons during the baseline period. Indeed, substantial fractions of granule cells showed excited or suppressed responses to orofacial movements during the baseline period (**Fig.4 E**, see *Methods*). Comparing the GLM movement weights which reflect the amplitude of movement- related responses, we found that movement-induced activities in granule cells, both excited and suppressed, became significantly weaker when mice engaged in the task (i.e. from baseline to ITI). In contrast, during passive exposure, this decrease was not observed; instead, the movement-excited neurons showed slightly enhanced movement-related responses (**Fig. 4F,G**).

The observation that task engagement increases odor responses and decreases movement responses of granule cells mirrors the pattern of cholinergic axons, which respond more strongly to odors and weakly to movements during task engagement (**Fig. 1I-K**). This raises the possibility that cholinergic modulation may contribute to the engagement-dependent modulation of the functional properties of granule cells. To test this possibility, we next examined the effect of inactivation of OB-projecting cholinergic neurons on granule cell activity. To minimize effects on animal behavior, we performed inactivation unilaterally by injecting retrograde AAV encoding Cre-dependent Flp and AAV encoding GCaMP6f into the dorsal OB, and AAV encoding Flp-dependent hM4D(Gi)-mCherry, or mCherry as a control, into the basal forebrain of ChAT-Cre mice (**Fig. 4H**). The animals were first trained to a 65-70% success rate. We then imaged granule cell activity for ∼70 trials (‘pre-CNO’), and then administered CNO intraperitoneally, and imaged the same group of granule cells for another ∼70 trials (‘post- CNO’).

As expected, this unilateral inhibition did not affect the animals’ behavior (**Fig. 4I**). Cholinergic inactivation, however, reduced odor responses of granule cells especially during task engagement. Only odor-suppressed responses were reduced during passive exposure. mCherry controls did not show these strong effects, even though there was a small but significant increase in odor-excited responses in both task and passive contexts, possibly reflecting CNO’s weak effect on the animal’s arousal state (**Fig. 4J**). In parallel, cholinergic inactivation decreased movement responses of granule cells during the baseline periods in both the task and passive conditions. Movement responses during the stimulus period (ITI) were also reduced in the passive condition but not during task engagement. The mCherry control group did not show such effects, confirming that the changes were due to the selective inactivation of cholinergic projection (**Fig. 4K**). These results support the idea that cholinergic axons in the OB enhance granule cell activity in a context-specific manner, selectively enhancing odor responses during task engagement and movement-related activity during other periods.

Finally, we examined the impact of OB-projecting cholinergic neurons on behavior by performing bilateral inactivation during training (**Fig. 4L**). To avoid tachyphylaxis from repeated CNO injections, we trained animals every two days and administered CNO in every training session. Over the 9 sessions of training, inactivation of OB-projecting cholinergic neurons led to significantly impaired performance compared to control mice expressing mCherry instead of hM4D(Gi), demonstrating the behavioral relevance of this population of cholinergic neurons.

After this period of repeated inactivation, hM4D(Gi)-expressing mice were trained without CNO, and they reached expertise within three training sessions, indicating that the deficits in these animals were caused by acute CNO injections (**Fig. 4M**).

In summary, these inactivation results suggest that cholinergic projections modulate OB activity in a context-specific manner, with their enhancement of odor responses during task engagement being crucial for precise odor-guided behaviors.

## Discussion

Here we revealed that OB-projecting cholinergic neurons show phasic odor responses selectively when mice are engaged in olfactory discrimination. The strength of these odor responses correlated with behavioral performance, and inactivation of these neurons impaired OB odor responses and behavioral performance. These findings align well with previous studies in rodents that showed the importance of acetylcholine in olfactory discrimination (*33, 34*). It is also noteworthy that the progressive loss of cholinergic neurons in Alzheimer’s and Parkinson’s diseases correlates with the progression of olfactory deficits, an early symptom in these neurodegenerative disorders (*35–38*). We propose that context-dependent modulation of olfactory processing by acetylcholine is critical for olfaction across species.

These odor responses dependent on task engagement were not seen in cholinergic neurons projecting to the dorsal cortex. These results suggest that cholinergic neurons operate through multiple parallel systems that modulate sensory processing in a modality-specific manner. It remains to be seen how many of these parallel systems exist. Is there a separate system for each sensory modality? Or are some sensory systems co-modulated? Furthermore, future studies are needed to determine the resolution of modulation—are they able to provide specific modulation within each modality to contribute to spatial or feature-specific attention? We found that OB- projecting cholinergic neurons project broadly across the olfactory pathway and also to other brain areas, but this does not preclude the existence of subpopulations that modulate the olfactory system with a finer resolution.

The brain has the remarkable ability to change its operation mode rapidly to adapt to ongoing behavioral demands. Providing a potential mechanism for this behavioral function, we found that functional tuning of cholinergic neurons can change rapidly when mice engage in a sensory task. The shift in tuning from movement to sensory may explain the recent finding of reduced movement responses during task engagement in neurons across cortical regions (*39*). This could also explain the dysfunctions in attentional shift in Alzheimer’s and Parkinson’s diseases that coincide with the loss of cholinergic neurons (*40, 41*). Rapid and context-dependent modulation of cholinergic neurons may be central to cognitive flexibility.

## Acknowledgments

We thank B. Morales, D. Arakelyan, A. Medina, and E. Hall for technical assistance; the rest of the members of the Komiyama lab, especially A. Wu and E. Gjoni, for discussions on the project and manuscript.

## Author contributions

Conceptualization: BY, TK Methodology: BY, TK Investigation: BY, YY Visualization: BY

Formal analysis: BY, RY Validation: BY

Resources: BY, CR, BL, TK Software: BY

Data curation: BY, YY Funding acquisition: TK Project administration: TK Supervision: TK

Writing – original draft: BY, TK Writing – review & editing: BY, TK

## Materials and Methods

### Animals

The experimental protocols were conducted in compliance with guidelines approved by the University of California San Diego’s Institutional Animal Care and Use Committee and NIH. ChAT-IRES-Cre knock-in mice (JAX 006410) were sourced from the Jackson laboratory. Adult mice, aged approximately 8-10 weeks, both males and females, were used for all surgical procedures and experiments. The animals were housed in standard plastic enclosures with typical bedding materials in a facility maintaining a reversed 12-hour light/dark cycle. Experimental activities were conducted exclusively during the dark phase. These mice were bred specifically for the studies detailed in this paper. No specific pairing of littermates or sexes was implemented for individual experiments. Throughout the training period, the mice’s health status was assessed daily.

### Viruses

The viral vectors were procured from Addgene, unless otherwise specified. For imaging cholinergic axon activity, we employed AAV5-hSyn-FLEX-axon-GCaMP6s (catalog number 112010) in ChAT-Cre mice. Functional imaging of granule cells (GCs) was achieved using AAV1-hSyn-GCaMP6f (catalog number 100837) in wildtype mice. In chemogenetic inactivation experiments, we utilized AAV8-hSyn-fDIO-hM4Di-mCherry (catalog number 154867) and AAVrg-phSyn1(s)-Flex-Flpo (catalog number 51669), while control animals received AAV8- Ef1a-DIO-mCherry (catalog number 114471), in ChAT-Cre mice. For anterograde tracing experiments, AAV1-CAG-fDIO-mNeonGreen (catalog number 99133) and AAVrg-phSyn1(s)- Flex-Flpo (catalog number 51669) were used in wildtype mice.

### Histology

For histological analysis, mice were anesthetized with a mixture of ketamine (150 mg/kg) and xylazine (12 mg/kg) administered based on body weight. They were then perfused transcardially with 4% paraformaldehyde. Following this, the brains were extracted and cryoprotected in a 30% sucrose solution until they sank. Using a Microm HM 430 microtome (Thermo Scientific), we cut the brains into 50 μm coronal sections. Finally, we mounted the slices using DAPI mounting medium (Sigma-Aldrich). Image acquisition of histological sections was performed using either an Olympus VS120 automated slide scanner or a Zeiss Apotome fluorescence microscope equipped with a 20× objective. To align the slide images with the Allen Brain Atlas, we employed two software tools: ABBA (Automated Brain Barrier Atlas) and QuPath. ABBA’s documentation can be accessed at https://abba-documentation.readthedocs.io/en/latest/, while information about QuPath is available at https://qupath.github.io/.

### Odorant delivery

Odorants used in all behavioral task and passive experience were diluted in mineral oil (Thermo Fisher Scientific, O121-1) to achieve a vapor pressure of 200 parts per million (ppm). Using a custom-built olfactometer, saturated odorant vapor was mixed with filtered, humidified air at a 1:1 ratio to a final concentration of 100 ppm. A mass flow controller (Aalborg, New York) was used to control the airflow at 1 L/min.

## Surgeries and viral injections

### Craniotomy

Anesthesia was administered to the mice using isoflurane, with an initial concentration of 3% for induction, followed by approximately 1% for maintenance. The skull was exposed, and a custom-designed stainless-steel headplate was affixed using superglue. A cranial window was created by performing a craniotomy over the right olfactory bulb. This opening was then covered with a glass window, which was sealed in place using vetbond and further secured with dental cement. Upon completion of the surgical procedure, the mice received subcutaneous injections of Baytril (10 mg/kg) and buprenorphine SR (0.5 mg/kg) for post-operative care.

### Injections

Horizontal puller equipment (P-2000, Sutter Instruments) was utilized to make glass pipettes from thin-walled glass capillary tubes (Wiretrol II). A custom-built injector was employed to administer viral solutions through these glass pipettes, which had an approximate diameter of 20 μm. The injection rate was maintained at about 20 nl/min.

Viral injections were performed stereotaxically prior to headplate implantation. In the case of basal forebrain injections, 600-700 nl of viral solution was administered per hemisphere over a 30∼35 minutes, injected at the following stereotaxic coordinates relative to bregma (in mm): 0.74 anterior, 0.65 lateral, 4.8 ventral. In the case of OB injection to label granule cells, the pipette tips were sharpened to a 50° to 60° angle using a custom rotation disk and viral solutions were injected, 20 nl over 3 min, at ∼0.4 mm beneath the dura surface. To minimize backflow, the pipettes were left in situ for an additional 15-20 minutes post-injection.

### Behavior

The behavioral protocols employed were largely consistent with our previous study (*42*). The water restriction regimen commenced two weeks post-surgery, approximately 15-18 days prior to the initiation of behavioral training. Daily water intake was limited to ∼1 ml to maintain the animals’ body weight at or above 80% of their initial weight. We utilized a real-time system (Rpbox, C. Brody) to manage the behavioral program. The experimental setup included two custom-fabricated lick ports, each equipped with infrared beams for detecting left and right licking actions. A trial was deemed correct or incorrect based on the initial lick during the designated answer period. A correct trial resulted in a water reward of ∼6 μl and an incorrect trial led to immediate trial termination without reward or punishment. Each mouse participated in one daily session, typically comprising 150 trials, unless the animal disengaged earlier.

### Pretraining

The behavioral training protocol was divided into three sequential stages. During the first stage, mice were incentivized to lick either the left or right lick port within a 2-second response interval for each trial, in the absence of any olfactory cues. The inter-trial interval (ITI) was gradually increased from 1 to 3 seconds. In the second stage, an odorant was introduced using cumene (Sigma-Aldrich, C87657) as the stimulus, presented for 4 seconds in each trial. Rewards were given for left port licking during the subsequent 2-second answer period. Licking of the right port during this time ended the trial without any punishment. The ITI was systematically extended from 3 to 15 seconds, increasing by 2-second increments, and then held constant at 15 seconds for the remainder of the training. The third stage employed 2-hexanone (Sigma-Aldrich, AC146881000) as the olfactory stimulus with a fixed 15-second ITI. In this phase, mice were rewarded specifically for right-side lick responses.

After achieving a performance accuracy exceeding 80% in these licking sessions, mice progressed to an easy discrimination task. This task involved the presentation of either cumene or 2-hexanone in each trial. The odorant on each trial was chosen randomly, with a maximum of three consecutive trials with the same odorant. Consistent with the licking sessions, cumene signaled left-lick trials, while 2-hexanone indicated right-lick trials. Odorant presentation was preceded by a 2 kHz, 2-second sound cue, delivered via a Piezo Buzzer Alarm. The odor valve opened 0.14 second after the sound offset. The odor reaches the animal 0.2 second after the odor valve open, which is estimated by the air flow speed and odor tube length. The ITI in each trial was a random integer between 13-17 seconds. Upon reaching a success rate of >80% with this initial set of easily distinguishable odorants (which took an average of 3.2 ± 1.7 sessions, mean ± SD), mice proceeded to a second easy discrimination task featuring a new odorant pair. In this subsequent task, cyclohexanone (Sigma-Aldrich, 398241) signaled left-lick trials, while isoamyl acetate (Sigma-Aldrich, C39601) indicated right-lick trials. Training with these new odorants continued until mice again achieved a >80% correct rate, typically requiring 1.9 ± 0.70 sessions.

### 4-odor discrimination task

In each trial, one of 4 binary mixtures [heptanal (Sigma-Aldrich, H2120) (%)/ethyl tiglate (Sigma-Aldrich, W246000) (%): H60E40, H53E47, H43E57 and H40E60] was delivered. Odorants with higher heptanal ratio signaled left-lick trials, and the odorants with higher ethyl tiglate ratio signaled right-lick trials. The rest of the settings were the same as the easy discrimination task. The mixture on each trial was chosen randomly with no more than three consecutive trials of odorants with the same lick side. Typically, the animals required 7 to 10 sessions of this task to consistently achieve over 80% correct responses. For the cholinergic axon recording group, 4 of the 6 mice were also trained to perform a second task. The second task has the same setting with the first one, but new odorants [binary mixtures of hexanal (Sigma-Aldrich, W255718) (%)/ethyl butyrate (Sigma-Aldrich, W242705) (%): H60E40, H53E47, H43E57 and H40E60] were used. Mice typically learned the second task more quickly, typically 5-7 sessions to consistently achieve over 80% correct responses.

### Passive experience

A separate group of mice underwent passive exposure without engaging in the task, but were otherwise treated identically to the task-performing animals, including water restriction. These mice were exposed to the same odorants within the same trial structure and session duration (150 trials), following the same behavioral timeline (pretraining, easy discrimination, and difficult discrimination). However, they did not participate in the actual discrimination task and no reward was given. The number of easy discrimination sessions for the passive experience group was determined using the median number of such sessions in the task engagement group. This resulted in three sessions of cumene versus 2-hexanone and two sessions for cyclohexanone versus isoamyl acetate. For 4-odor passive exposure, only the binary mixtures of heptanal and ethyl tiglate were used.

### Cholinergic axonal functional imaging

Functional imaging of cholinergic axons was conducted during behavioral sessions of the 4-odor discrimination task (or 4-odor passive exposure) at regular intervals, generally every three days to minimize photobleaching and phototoxicity. The imaging was typically performed on Day 1, Day 4, Day 7, and Day 10. However, if mice attained expert performance (success rate >80%) before Day 10, the imaging schedule was modified accordingly. For the group undergoing passive experience, imaging was done every three days, mostly on Day 1, Day 4, and Day 7. Across imaging days, the same fields were consistently imaged, with typically two imaging fields recorded in each session.

### Somatic functional imaging

For functional somatic imaging of granule cells, imaging was conducted in the middle of the 4- odor task or passive exposure, typically between days 4-6. For the task animals, the exact day of imaging was selected once mice reached a success rate of at least 60% and showed no clear bias toward one side. The imaging fields were carefully matched before and after CNO administration to ensure consistency in data collection.

### Chemogenetic manipulation

ChAT-Cre mice expressing hSyn-fDIO-hM4Di-mCherry in the basal forebrain cholinergic neurons underwent behavioral training as previously described. Clozapine-N-oxide (CNO, Enzo Life Sciences) was prepared by dissolving it in deionized water to a concentration of 2.5 mg/mL. For the imaging experiment, a field of granule cells was imaged for 70-75 trials. Subsequently, CNO was administered intraperitoneally at a dose of 8 mg/kg body weight. Immediately after the injection, the mice were returned to their home cages. After 30-minutes, the same group of granule cells was imaged for an additional 70-75 trials. For repeated inactivation throughout task training, CNO was injected intraperitoneally at a dose of 5 mg/kg body weight 30 minutes prior to behavioral training. The training procedures were identical for both the manipulation and control groups. Littermates from these two groups were trained in parallel on the same days.

### Image acquisition

*In vivo* imaging was conducted using a commercial two-photon microscope (B-scope, Thorlabs) with 925-nm excitation (Mai Tai, Spectra-physics) at a frame rate of approximately 30 Hz. For soma imaging, each frame covered a field of view of about 308 × 264 μm, while for cholinergic axon imaging, the field was approximately 142 × 121 μm, both with a resolution of 512 × 512 pixels. To ensure consistency across sessions, the average image from the first imaging session was used as a template for subsequent sessions. Frame times were recorded and synchronized with behavioral recordings using Ephus software. Manual corrections were made during imaging to address slow drifts in the field of view, using reference images. For cholinergic axon imaging in the dorsal cortex, three cortical areas were investigated: primary somatosensory cortex (S1, centered at 1.8 mm lateral and 0.75 mm posterior to bregma), primary visual cortex (V1, centered at 2.5 mm lateral and 3.25 mm posterior to bregma), and primary motor cortex (M1, centered at 1.5 mm lateral and 0.4 mm anterior to bregma).

## Data analysis and statistics

### Regions of interest (ROI) detection and fluorescence analysis

Time series images were initially saved as Tiff files and processed using suite2P (*43*) for motion correction. For both cholinergic axons and granule cells, ROIs were first identified using suite2P and then refined through visual inspection. ROIs clearly belonging to the same axon were manually combined. For cholinergic axons, in each field of view, three background ROIs were manually selected from dark areas without obvious fluorescent transients, and the average fluorescence time series of these background ROIs served as the background for all ROIs within the same field of view. For granule cells, the background neuropil fluorescence is estimated using the built-in function in suite2P. For fluorescence analysis of cholinergic axons, the pixel values within each ROI were averaged, and the background fluorescence time series was subtracted. For granule cells, the corrected fluorescence is F (the ROI fluorescence) – 0.7 × Fneu (neuropil fluorescence). The raw fluorescence signals were then filtered using a Savitzky-Golay filter with 1-second window for granule cells and 2-second window for cholinergic axons and 3 for the polynomial order. Baseline estimation was performed using the OASIS package (https://github.com/j-friedrich/OASIS).

### Generalized linear model

To assess the relationship between facial movement and sensory stimuli and neural activity, two generalized linear models were used. All behavioral variables were downsampled to 30 Hz to align with the imaging frame rate. One model utilized only the movement (analog) predictor, while the other employed sensory stimulus (digital) predictors to forecast cholinergic axon activity.

For the analog movement GLM, we used:

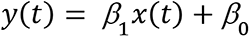

Where y(t) represents the averaged z-scored calcium time series of all cholinergic axons within the imaging field during the baseline and inter-trial interval (ITI). The time series was z-scored across the entire session. x(t) denotes the orofacial motion energy as the single movement predictor during the same period. x(t) was time-shifted based on the peak of cross-correlation between the calcium and motion energy, typically by 3 frames (∼100ms). The ITI was defined as the period between 3.5 seconds after the end of the answer period and one second before the sound cue of the next trial and these ITIs were concatenated to generate a single time series.

Since the ITI time series was longer than the baseline, for fair comparison, a consecutive segment of ITI matching the length of the baseline was selected to perform the regression. The selection of ITI segments was repeated 50 times and the results were averaged.

For the digital sensory stimulus GLM, we employed:

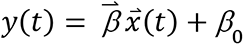

Here, y(t) is the averaged z-scored calcium time series of all cholinergic axons within the imaging field throughout the entire stimulus period. 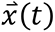 represents the sensory stimuli, including odorant and sound predictors. Odorant predictors consisted of four 1-sec digital pulses for each second during the odor period, and an additional 0.5-second pulse after odor offset.

Sound predictors comprised three digital pulses per second, and a 0.3-second pulse after sound offset. All of the 8 predictors are linearly independent with each other. To mitigate overfitting, ridge regression was applied to both models. The performance of both models was evaluated using ten-fold cross-validated correlations between predicted and actual imaged activity, yielding R^2^ values. The odor and sound weights shown in Fig.1 and Fig. 3 are the average weights of all predictors for the odor and sound, respectively.

### Decoding odor identity

A support vector machine with radial basis function kernel was used to decode the odor identity with scikit-learn package in Python. Cholinergic axonal response in each trial was expressed as a population activity vector by averaging ΔF/F0 values 0-4 s of the odorant period for each axon segment. For each session, 80% of trials were used for training the decoder and 20% were used for validation (5-fold validation), and this was repeated 300 iterations with a random selection of training trials for each iteration. The decoding accuracy of each session is the average of the 300 iterations, and the fraction of the iterations that had the decoding accuracy below the chance level (25%) defined the p value for decoding significance.

### Classification of movement-related neurons

To identify movement-related neurons, the movement GLM described previously was applied to each granule cell’s calcium time series during the baseline period of the recording before CNO injection. A neuron was classified as movement-related if it met two criteria: *1.)* the movement predictor showed statistical significance (P < 0.01), and *2.)* the variance explained by the model exceeded 1% (R^2^ > 0.01).

### Defining responsive odor-ROI pairs

Responsive odor-ROI pairs were defined as previously reported (*42*). An odor-ROI pair was classified as responsive if the following two criteria were met:

1. Within any sliding window of 0.5 s (15 frames) during the odor period, there are more than 75% of the image frames where *F*/*F*0 of all trials are significantly different (Wilcoxon rank-sum test, *P* < 0.01) from the baseline frames (average of the 5-s period before sound onset) of all trials.
2. The average *F*/*F*0 of all trials in at least one frame during the 0.5-s window that meets criterion 1 is at least 0.2 larger (excited) or smaller (suppressed) than average baseline fluorescence of all trials.

### GLM-HMM model

A 3-state hidden Markov model with Bernoulli generalized linear model observations (GLM-HMM) (*28*) was used to classify trials into two categories: engaged and biased. A universal model was trained using selected sessions to balance different conditions and was then applied to all task sessions, regardless of behavioral performance. The model is defined by a KxK transition matrix (where K represents the number of states) and a set of weights for each state. Three input parameters were used: input stimulus, choice, and previous choice. The model was trained using the expectation-maximization algorithm. The initial weight setting was critical for the model performance, and to identify the optimal initial weights, five sessions exemplifying obvious fully biased, fully engaged, and mixed states were manually labeled as ground truth. 50 different sets of initial weights were tested and the one that best predicted the ground truth was used for analysis. The primary code workflow adhered to the methodology outlined in a previous study (*28*).

## Statistical analysis

All statistical analyses were performed with Python using scipy or statsmodels package. Standard statistical tests used are described in figure legends.

For the mixed effects model that compared movement variance or weights between the ITI and baseline phase for dorsal cortex cholinergic axon recording in Fig 3, we used the model below:

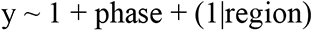

where the fixed effect is the phase, baseline or ITI, and a random intercept for each region.

For the model that compared movement weights between baseline and ITI:

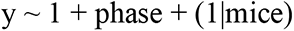

where the fixed effect is the phase, which is ITI or baseline, with a random intercept for each mouse.

For the model that compared the odor response or Δmovement weight between task and passive conditions:

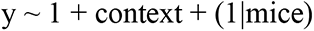

where the fixed effect is the context, which is either ITI or baseline, with a random intercept for each mouse.

For the model that compared the baseline and ITI movement weights and odor responses between pre- and post-CNO administration:

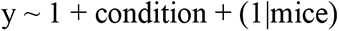

where the fixed effect is the condition, which is either pre- or post-CNO, with a random intercept for each mouse.

---

**Fig. S1.**
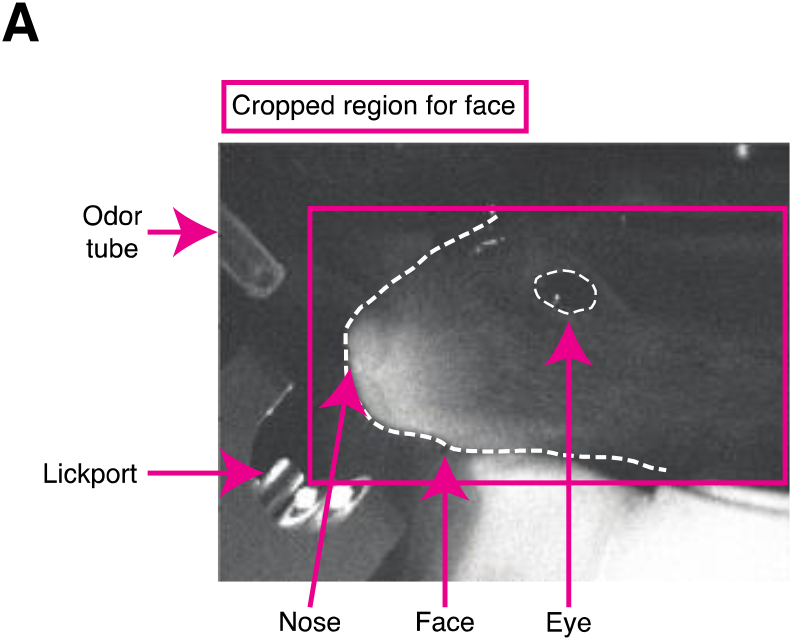
Example video frame for orofacial motion analysis. (A) Representative video frame captured by the camera during the experiment. The purple rectangle indicates the region of interest (ROI) used to compute orofacial motion energy.

